# Transposable element competition shapes the deer mouse genome

**DOI:** 10.1101/2022.10.18.512801

**Authors:** Landen Gozashti, Cedric Feschotte, Hopi E. Hoekstra

**Affiliations:** Department of Organismic & Evolutionary Biology, Department of Molecular & Cellular Biology, Museum of Comparative Zoology and Howard Hughes Medical Institute, Harvard University, 16 Divinity Avenue, Cambridge, Massachusetts 02138, USA; Department of Molecular Biology & Genetics, Cornell University, Ithaca, New York 14850, USA

## Abstract

The genomic landscape of transposable elements (TEs) varies dramatically across species, with some TEs demonstrating greater success in colonizing particular lineages than others. In mammals, LINE retrotransposons typically occupy more of the genome than any other TE and most LINE content is represented by a single family: L1. Here, we report an unusual genomic landscape of TEs in the deer mouse, *Peromyscus maniculatus*, a model for studying the genomic basis of adaptation. In contrast to other previously examined mammalian species, LTR elements occupy more of the deer mouse genome than LINEs (11% and 10% respectively). This pattern reflects a combination of relatively low LINE activity in addition to a massive invasion of lineage-specific endogenous retroviruses (ERVs). Deer mouse ERVs exhibit diverse origins spanning the retroviral phylogeny suggesting that these rodents have been host to a wide range of exogenous retroviruses. Notably, we were able to trace the origin of one ERV lineage, which arose within the last ∼11-18 million years, to a close relative of feline leukemia virus, revealing inter-ordinal horizontal transmission of these zoonotic viruses. Several lineage-specific ERV subfamilies have attained very high copy numbers, with the top five most abundant accounting for ∼2% of the genome. Concomitant to the expansive diversification of ERVs, we also observe a massive expansion of Kruppel-associated box domain-containing zinc finger genes (KZNFs), which likely control ERV activity and whose expansion may have been partially facilitated by ectopic recombination between ERVs. We also find evidence that ERVs directly impacted the evolutionary trajectory of LINEs by outcompeting them for genomic sites and frequently disrupting autonomous LINE copies. Together, our results illuminate the genomic ecology that shaped the deer mouse genome’s TE landscape, opening up a range of opportunities to investigate the evolutionary processes that give rise to variation in mammalian genome structure.

**Summary:** Transposable elements (TEs) are a highly diverse collection of genetic elements capable of mobilizing in genomes and function as important drivers of genome evolution. The landscape of TEs in a genome have been compared to a genomic ecosystem, with interactions between TEs and each other as well as TEs and their host, dictating the evolutionary success of TE lineages. While TE diversity and copy numbers can vary dramatically across taxa, the evolutionary reasons for this variation remain poorly understood. In mammals, long interspersed nuclear elements (LINEs) typically dominate, occupying more of the genome than any other TE. Here, we report a unique case in the deer mouse (*Peromyscus maniculatus*) in which long terminal repeat (LTR) retrotransposons occupy more of the genome than LINEs. We investigate the evolutionary origins and implications of the deer mouse’s distinct genomic landscape, revealing ecological processes that helped shape its evolution. Together, our results provide much-needed insight into the evolutionary processes that give rise to variation in mammalian genome structure.

## Introduction

Transposable elements (TEs) are parasitic genetic elements capable of mobilizing in genomes and function as important drivers of genome evolution [1–3]. In mammals, for example, TEs account for at least 20% of the genome and, in some cases, have been exapted for significant functional innovations [4–6]. Nonetheless, TEs generate mutations when they insert into new positions in the genome and thus can represent a significant burden on host fitness. This cost is compounded by the fact that TEs can contain gene regulatory sequences and cause structural rearrangements even after they have lost the ability to transpose [1,7]. The evolutionary success of a given TE lineage is dictated by its ability to replicate faster than the host genome but limited by its cost to host fitness [3,8]. TE lineages are in a constant coevolutionary conflict with each other as well as their host [9,10]. As a consequence, hosts have evolved various ways to suppress TE activity [11]. These genetic conflicts embody the “ecology of the genome” and play an important role in shaping the genomic landscape of TEs in a given species as well as its genome structure more broadly [9,10].

TEs are remarkably diverse, and TE landscapes can vary dramatically across species [12]. TEs are classified into two broad categories based on their transposition mechanism: class I elements (retrotransposons), which mobilize through an RNA intermediate, and class II elements (DNA transposons), which do not. Most eukaryotic lineages harbor a diversity of TEs from multiple taxonomic subgroups within each of these broad classes [1,12]. By contrast, some phylogenetic groups have TE landscapes that are relatively similar across species [13,14]. One such clade is mammals [5]. In most mammalian genomes, DNA transposons cannot actively mobilize and only exist as relics of anciently active elements [5]. Actively mobilizing retrotransposons include long terminal repeat retrotransposons (LTR), which are mostly endogenous retroviruses (ERVs), as well as non-LTR retrotransposons represented by long interspersed nuclear elements (LINEs) and their nonautonomous counterparts, short interspersed nuclear elements (SINEs) [5,15]. LINEs are nearly always the most abundant TEs, and most are represented by a single family, L1, which typically occupies hundreds of megabases of the mammalian genome [5]. However, the dearth of examples of alternative TE landscapes has limited our ability to investigate the evolutionary processes driving mammalian genome structure evolution and specifically, the maintenance of LINE dominance [5,16].

The North American deer mouse, *Peromyscus maniculatus*, has become an important model for studying the genetic basis of adaptation [17]. Early studies of deer mice and closely related species used polymerase chain reaction (PCR) methods to explore TE abundance and reported evidence for an unprecedented expansion of endogenous retroviruses (ERVs) [18,19]. However, the landscape of TEs in the deer mouse remains unexplored on a genomic scale. Here, we report a highly distinct genomic landscape of TEs in the deer mouse genome. We find that, in contrast to nearly all examined mammalian genomes, LTR retrotransposons are more abundant in the deer mouse genome than LINEs. We investigate the evolutionary origins and implications of the deer mouse’s distinct genomic landscape, revealing ecological processes that helped shape its evolution.

## Results and Discussion

### Deer mice exhibit a unique landscape of transposable elements

To evaluate the genomic landscape of TEs in the deer mouse genome, we first generated a lineage-specific TE library *de novo* from the deer mouse (*P. maniculatus bairdii*) genome using a combination of systematic and manual methods (see Methods). We identified 48 LINE, 28 SINE, and 118 LTR deer-mouse specific subfamilies (Figure 1A; Supplementary Table 1). We then merged this lineage-specific TE library with all curated mammalian TEs from the Dfam database [20] and annotated the genome using the combined library. We define lineage-specific subfamilies with respect to those observed in house mice, *Mus musculus* (strain C57BL6). Our annotation revealed a distinct genomic landscape of TEs in the deer mouse, relative to other mammals, in which LTR elements occupy more of the genome than LINEs (Figure 1A). Specifically, LTR elements occupy ∼11% of the genome, followed by LINEs (∼10%), SINEs (7%), and other TEs (<2%) (Figure 1A; Supplementary Table 2). It is also worth noting that the dearth of LINE content observed in the deer mouse genome is unlikely an artifact of our inability to detect lineage-specific LINEs since vertical propagation of LINEs has been accompanied by relatively little sequence changes. In total, TEs occupy ∼30% of the deer mouse genome, reflecting an increase in TE content relative to other species in the rodent Family Cricetidae, such as the grasshopper mouse (*Onychomys torridus*, 24%) and prairie vole (*Microtus ochrogaster*, 17%) (Figure 1A), but a considerable reduction relative to house mouse (*Mus musculus*, >40%), although these differences may reflect, at least in part, differences in genome assembly and TE annotation quality [21,22]. Nonetheless, most of the difference in TE content between the deer mouse and house mouse can be attributed to decreased LINE content in the deer mouse, whereas most of the difference in TE content among cricetid species can be attributed to LTR elements.

**Figure 1:**
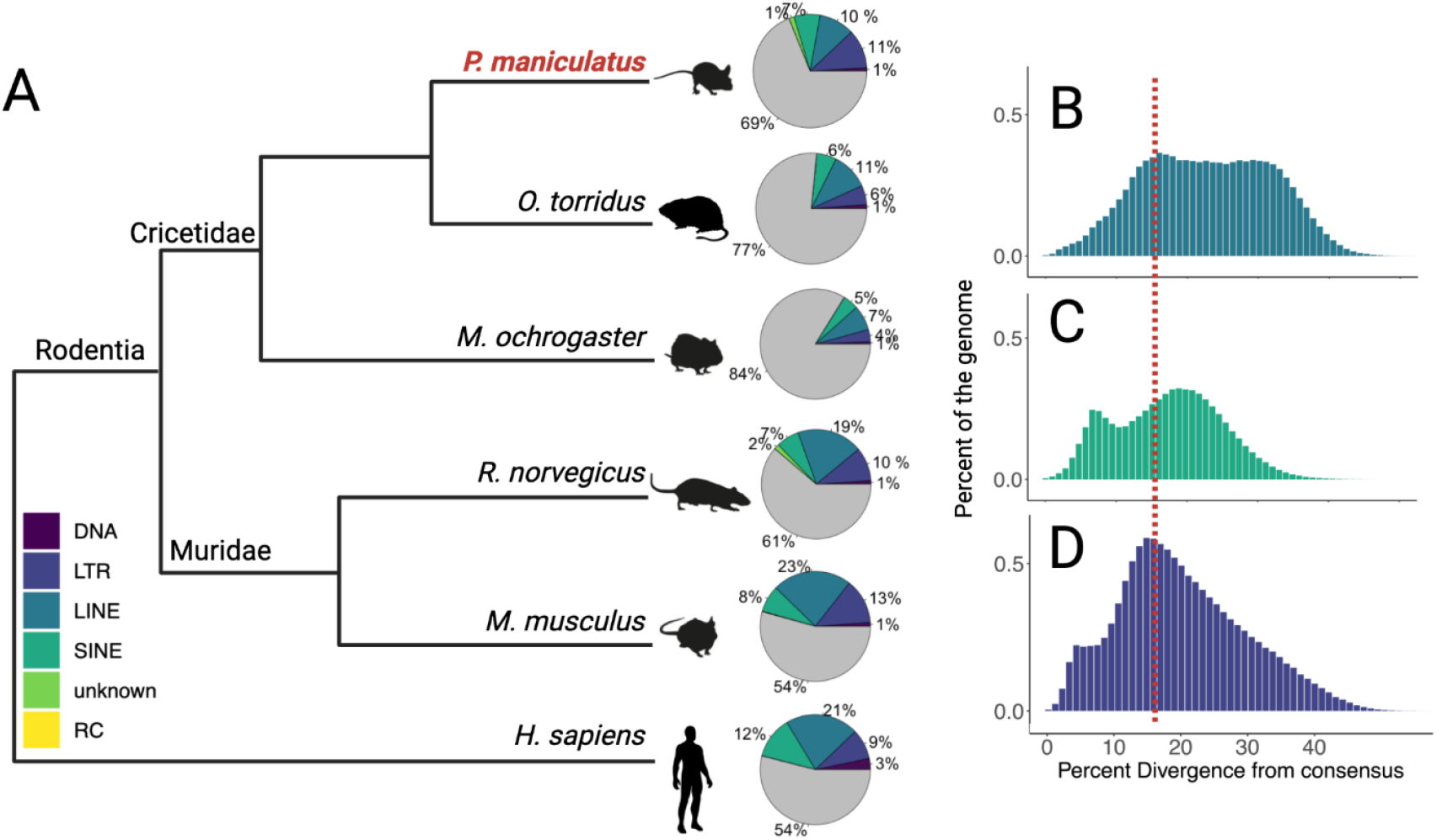
Transposable element landscape. (**A**) Cladogram highlighting the relationship of deer mice (*P. maniculatus*; red) to other mammalian species considered in this study. Pie charts show the relative percent of the genome occupied by TE subclasses for each species. Color corresponds to the percent of the genome attributed to each type of TE (see legend); gray represents the percent of the genome that is not occupied by TEs. Note, in *P. maniculatus*, LTRs (dark blue) occupy more of the genome than LINEs (aqua blue). (**B-D**) Percent of the genome as a function of CpG corrected Kimura divergence from the consensus for each TE subfamily of (**B**) LINEs, (**C**) SINEs, and (**D**) LTR elements. The red dotted line represents the start of observed LINE decline in all plots.

Based on these observations, we hypothesized that the unique TE landscape of deer mice is the result of a combination of reduced lineage-specific LINE gain and a rapid proliferation of lineage-specific LTR elements. To investigate this possibility, we first compared genomic representation as a function of within-subfamily divergence across LINEs, SINEs, and LTR elements (Figure 1B-D). Consistent with our hypothesis, we observe reduced representation of LINEs with lower divergence from the consensus, suggesting decreased LINE accumulation in the deer mouse lineage on more recent timescales (Figure 1B). However, despite this decline in the accumulation of LINEs, we still find multiple candidate LINEs with intact protein machinery, suggesting that LINEs are still active, consistent with previous reports of LINE activity in deer mice (Supplementary Table 3) [23]. We also observed evidence for lineage-specific SINE activity (Figure 1C). Since SINEs parasitize LINE machinery for mobilization, evidence of recently active SINEs suggests that potentially mobile LINEs still exist in the genome. In addition, we find a recent lineage-specific proliferation of LTR elements (Figure 1D): LTR elements are significantly overrepresented among the youngest TEs in the genome (<1% divergence from the consensus; two-sided Fisher’s exact test, P<0.00001). Furthermore, the observed decline of LINE gains in the genome coincides with the peak of LTR gains in the genome (Figure 1B-C). Together, these results suggest that both reduced LINE gain and lineage-specific LTR proliferation have contributed to the deer mouse’s unique TE landscape, and that the two may be associated.

### DNA loss fails to explain reduced LINE content

In addition to gain, TE loss can be an important driver of genomic TE content. Although we find evidence for a decline of LINE gain, the low LINE content in the deer mouse genome, relative to house mouse, could also have resulted from higher rates of loss in the deer mouse (Figure 2A-B). To investigate this possibility, we calculated the DNA loss coefficient *k* (following [24]), using the formula *E = A e-kt*, where *E* is the amount of extant ancestral DNA in the species considered, *A* is the ancestral assembly size, and *t* is time. Larger values of *k* suggest higher rates of lineage-specific DNA loss. We calculated a *k* coefficient of ∼0.0047 for deer mice, a value similar to, and in fact slightly lower than the k value estimated for house mouse (∼0.006), suggesting that the reduced LINE content observed in the deer mouse genome cannot be explained by significantly higher rates of loss in deer mice (Figure 2C).

**Figure 2:**
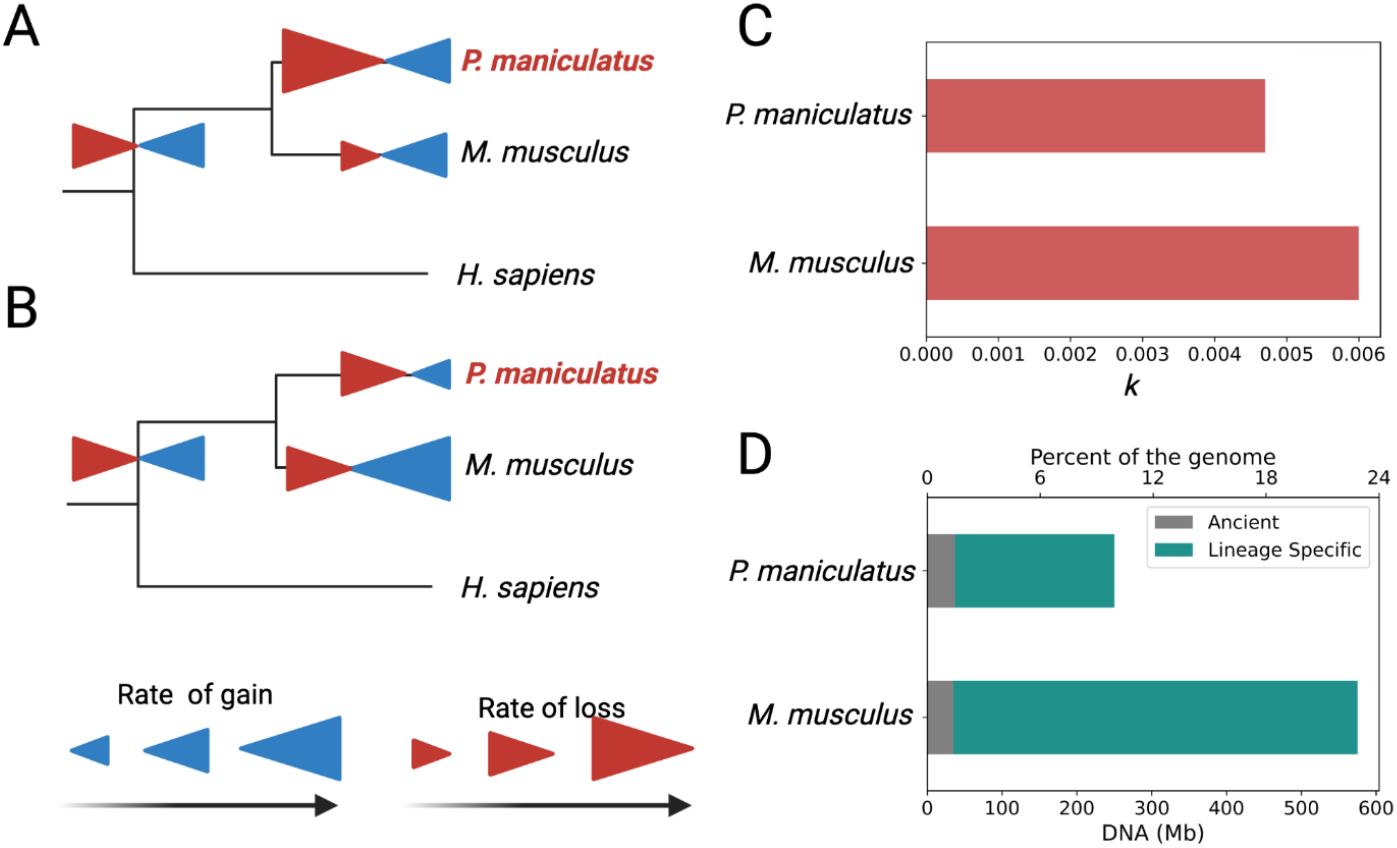
(**A**-**B**) Non mutually exclusive evolutionary scenarios that may have shaped the TE landscape in the deer mouse. (**A**) High rates of lineage-specific loss could have resulted in the reduced LINE content observed in deer mouse (*P. maniculatus*) relative to house mouse (*M. musculus*), in addition (**B**) to differences in lineage-specific TE activity. (**C**) *k* coefficients of DNA loss for *P. maniculatus* and *M. musculus* suggest lower rates of loss in *P. maniculatus*. (**D**) Genomic representation of ancient (gray) and lineage-specific (green) LINEs in *P. maniculatus* and *M. musculus* reflect greater representation of ancient elements in *P. maniculatus* relative to *M. musculus*.

Nonetheless, it remains possible that the rate of LINE loss in the deer mouse does not reflect the genome-wide rate of loss; rather, LINEs are lost at a higher rate. To investigate this possibility, we compared the proportions of DNA attributed to ancient mammalian LINEs present in the common ancestor of the deer mouse and house mouse as well as lineage-specific elements. If the relative absence of LINEs in the deer mouse is due to high rates of loss, we expect to find a decreased amount of DNA attributed to ancient LINEs in the deer mouse relative to house mouse. To the contrary, we find that although LINEs contribute to over twice as much of the house mouse genome as the deer mouse genome (∼575Mb as opposed to ∼250Mb), ancient LINEs are significantly less represented in the house mouse genome (∼8% of total LINEs versus 2% in the deer mouse; two-way Fisher’s Exact Test, P=0.006; Figure 2D). These results suggest that the low LINE content observed in the deer mouse cannot be explained by high rates of LINE-specific loss, and instead, is more likely the result of reduced rates of gain.

### Nonautonomous ERVK-like elements are abundant

Most mammalian LTR retrotransposons are endogenous retroviruses (ERVs). ERVs are divided into three broad classes depending on their retroviral origins: ERV1, ERVK and ERVL, with an additional subgroup of nonautonomous ERVL-MaLRs [20,25]. In the deer mouse, we find a pattern in which ERVK and ERVL-MaLR elements together account for over 80% of genomic ERV content, consistent with previous observations in other rodents [5,20] (Figure 3A). Lineage-specific elements represent over half of genomic ERV content (∼57%), further suggesting that the deer mouse has experienced a lineage-specific ERV expansion. However, when we compare the relative genomic proportion of ERVs across lineage-specific subfamilies, we find that ERVKs account for a disproportionately large proportion of ERV content relative to shared elements (Fisher’s exact test, P<0.00001), representing over 75% of observed lineage-specific ERV sequence in the genome (Figure 3B; Supplementary Table 1).

**Figure 3.**
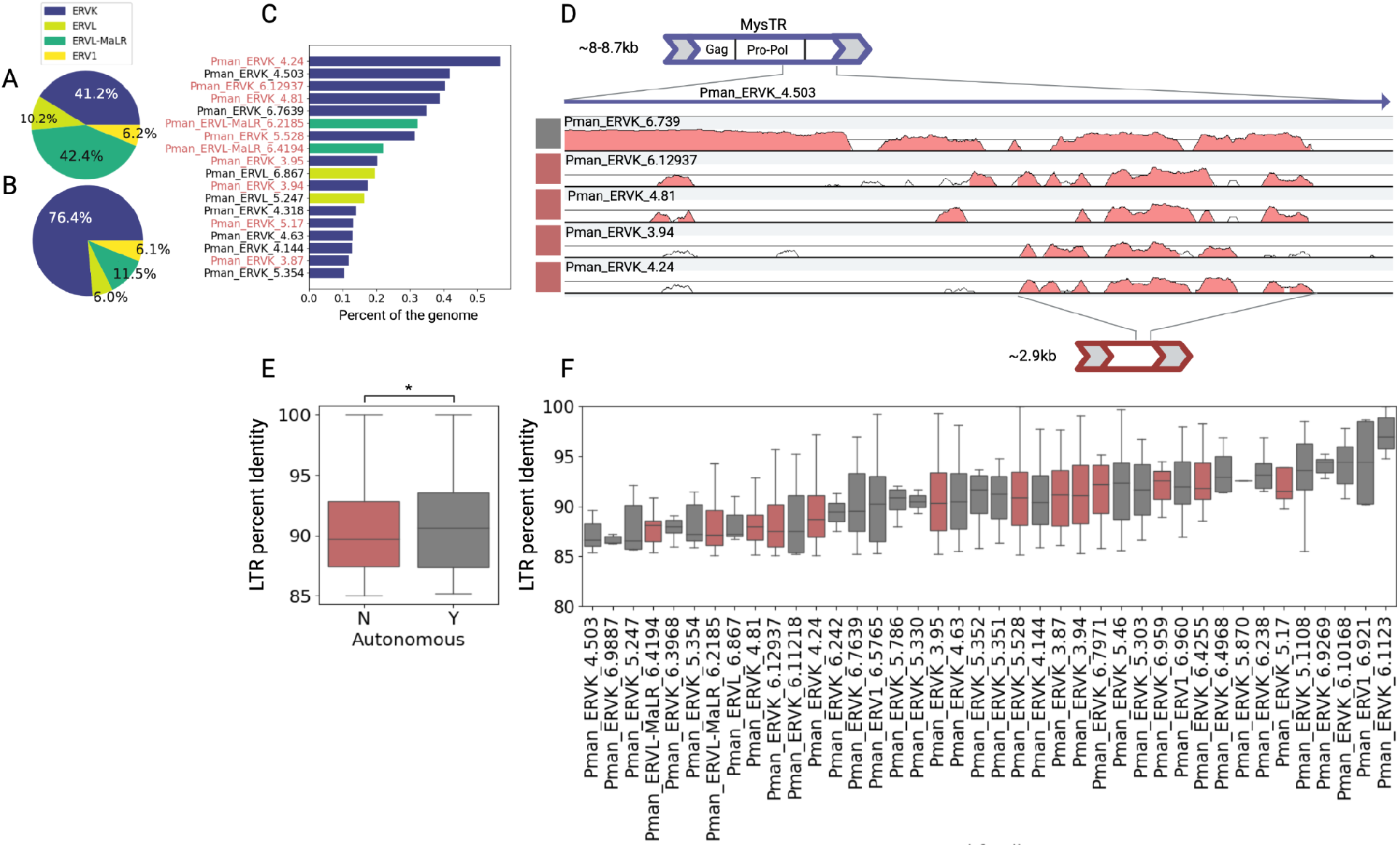
Relative contribution of different broad ERV classes to ERV content in the deer mouse across (**A**) all ERVs and (**B**) lineage-specific ERVs. (**C**) Respective genomic occupancy across the 18 most common lineage-specific ERV subfamilies. Red font denotes nonautonomous subfamilies. (**D**) VISTA plot showing regions of homology between autonomous mysTR subfamilies and nonautonomous subfamilies as well as their respective position. Gray boxes on the left denote autonomous subfamilies, and red boxes denote nonautonomous ones. (**E**) Comparison LTR percent identity aggregated across all lineage-specific autonomous and nonautonomous elements suggests that autonomous elements are significantly younger (Man Whitney U, *p<0.05). (**F**) Comparison of LTR percent identity across all ERV subfamilies that display at least one candidate full-length copy showing that the youngest subfamilies are autonomous (gray).

ERVKs are typically autonomous elements that encode their own machinery for mobilization [26]. However, we manually inspected deer mouse*-*specific ERV subfamilies, annotating *gag, pro, pol* and *env* genes as well as the protein domains required for autonomous transposition. We found that several of the most abundant ERVK families did not possess any internal open reading frames (ORFs) or expected protein domains, suggesting that they might be nonautonomous. Many ERVs contained gaps that interrupted or truncated their internal sequences (507 of 1119 candidate full length ERVs), making it challenging to reconstruct full length elements and assess the presence or absence of coding machinery. In light of this, we required that a putatively nonautonomous ERV subfamily display at least five full length copies with no gaps for it to be classified as nonautonomous, regardless of its consensus sequence length or content. Even with this conservative filter, we found that the most abundant deer mouse-specific ERV subfamilies are nonautonomous ERVK-like elements (Figure 3C; Supplementary Table 4).

Pman_ERV2_4.24, for example, is the most abundant ERV in the genome, accounting for ∼5% of total ERV content. Furthermore, for the subset of ERVKs in which we could confidently reconstruct full length sequences and assess autonomy, nonautonomous elements (53%) occupy significantly more of the genome than autonomous ones (47%; Fisher’s exact test p<0.0001; Supplementary Table 4). Overall, our results suggest that ERVKs and their nonautonomous counterparts have had a significant impact on the deer mouse’s unique genome structure.

### Several ERVKs show sequence homology to mysTR

Nonautonomous TEs parasitize autonomous elements for mobilization. Studies on the nonautonomous ERVK, ETn, in house mouse showed that ETn exhibits regions of homology to fully coding MusD ERVKs, suggesting that ETn likely arose from the ancestors of Mus-D [27] and now hijacks MusD machinery for mobilization via complementation in trans [28]. Given the massive copy numbers of nonautonomous ERVKs in the deer mouse genome, we next searched for related autonomous elements that may facilitate nonautonomous element mobilization. In addition to several prolific nonautonomous subfamilies, we observe three autonomous ERVK subfamilies, Pman_ERVK_4.503, Pman_ERVK_6.7639 and Pman_ERVK_5.247, that together occupy a remarkable 1% of the genome (Figure 3C). These subfamilies show sequence homology to mysTR, a previously identified ERV [18].

Previous studies failed to identify full-length mysTR copies with intact *gag* and *pro-pol* genes required for mobilization, raising questions about mysTR’s overall autonomy and current ability to mobilize, although these studies lacked the genomic resources to analyze mysTR sequences on a genome-wide scale [18,29]. Our genome-wide analysis reveals multiple candidate intact copies of mysTR related ERVs, which display ORFs with homology to *gag* and *pro-pol* genes and contain expected protein domains, suggesting that these ERVs are indeed autonomous and still active in the deer mouse (Supplementary Table 4). Furthermore, we hypothesized that observed nonautonomous ERVKs represent mys elements, which are also related to mysTR and may now use mysTR protein machinery for mobilization [19,30]. Indeed, similarly to ETn and MusD, we find regions of homology between autonomous mysTR related ERVK subfamilies and nonautonomous ERVK subfamilies (Figure 3D; Supplementary Table 4). The most conserved region of nucleotide sequence homology between these subfamilies is just downstream of the *pro-pol* gene and upstream of the 3’ LTR (Figure 3D). Interestingly, candidate nonautonomous and autonomous mysTR-related subfamilies do not display strong homology outside of this region, suggesting that ERVKLs may have evolved through an internal recombination event in an autonomous ERVK [27]. Maintenance of sequence similarity in this region is also consistent with strong selection due to a possible role in ERVK mobilization, although its function remains unknown.

### Autonomous ERV subfamilies are younger than autonomous ones

Since ERV LTRs are identical upon insertion, LTR sequence identity can provide an estimate for how recently an ERV inserted. To investigate the evolutionary dynamics of deer mouse ERVs, we compared the distributions of LTR identity across lineage-specific ERV subfamilies. We find that nonautonomous ERVs are significantly older than autonomous ERVs (Mann Whitney U, P=0.03; Figure 3E). This pattern is likely explained by multiple non-mutually exclusive processes. First, nonautonomous ERV insertions likely rise to fixation more often than fully autonomous insertions since they are less deleterious to their host. This is expected since nonautonomous ERVs are shorter than their autonomous counterparts and lack coding transcripts which can be toxic to the host. Second, since the coding capacity of autonomous ERVs dictates the ability of both autonomous and nonautonomous ERVs to mobilize, the activity of parasitic nonautonomous subfamilies is likely associated and limited by that of their autonomous targets [31]. Consistent with this, we also observe that the youngest ERV subfamilies in the deer mouse genome are autonomous (Figure 3F).

### Diverse origins of endogenous retroviruses

ERVs in the deer mouse genome arose from diverse retroviruses ERVs arise in a species when an exogenous retrovirus infects the germline, and new families of ERVs evolve *de novo* more frequently than other autonomous mammalian TEs such as LINEs [26]. To investigate the retroviral origins of ERVs in the deer mouse, we focused on full-length ERVs across all identified subfamilies with flanking LTRs, *pol* genes and full-length reverse transcriptase (RT) domains (>450bp), which we used for classification and phylogenetic analysis. We initially identified 148 candidate full-length ERVs with *pol* genes and evidence of an RT domain. However, many ERVs contained ambiguous sites or gaps that interrupted or truncated the RT domain, leaving only 52 ERVs which met our conservative requirements (Supplementary Table 5-6). Thus, we note that our reported ERV diversity is likely an underestimate.

We initially used a hidden Markov model (HMM) approach [32] to classify ERVs based on their RT domains (Supplementary Table 6). Using this approach, we find that of the 52 deer mouse ERVs with full-length RT domains, 11 are derived from gammaretroviruses (ERV1), 39 from betaretroviruses (ERVK), and 2 from spumaretroviruses (ERVL) (Supplementary Table 6). Phylogenetic analysis of RT domains from these ERVs and other known retroviruses supports these initial classifications and shows that deer mouse ERVs form 14 distinct clusters representing at least 14 independent origins spanning retroviral diversity (Figure 4A). Most of these are derived from diverse betaretroviruses (9 of the 14), consistent with previous observations in other rodents [33,34]. Additionally, four ERV clusters show evidence of gammaretroviral origin, and one ERV cluster shows evidence of spumaretroviral origin (Figure 4A).

**Figure 4:**
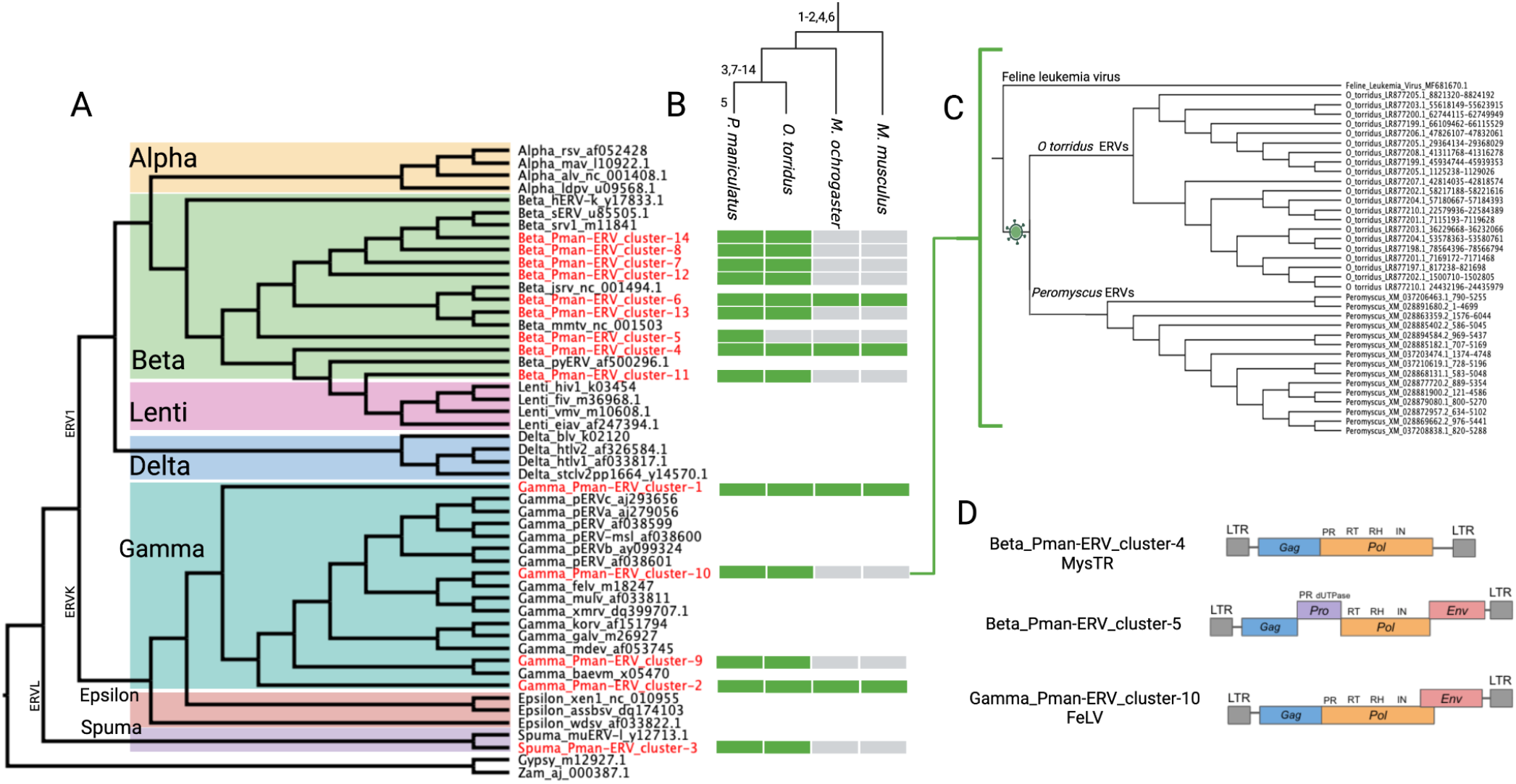
Origins of ERVs in the deer mouse genome. (**A**) Phylogeny constructed with *RT* domains of deer mouse ERVs and publicly available endogenous and exogenous retroviruses for context. Deer mouse ERVs form 14 distinct clusters spanning broad retroviral diversity. (**B**) Cladogram showing the approximate time of origin of deer mouse (*P. maniculatus*) ERVs based on presence or absence in three other species at different phylogenetic distances. ERV cluster numbers are annotated on the branch corresponding to their approximate origin. For each species, green boxes represent presence and gray boxes represent absence for each respective ERV cluster found in *P. maniculatus*. (**C**) Neighbor-joining tree showing the phylogenetic relationship between Pman-ERV_cluster-10 copies in *P. maniculatus*, related ERVs in grasshopper mouse (*O. torridus*), and feline leukemia virus. (**D**) Structure of ERVs found in *P. maniculatus* with ORFs denoted by colored boxes and important protein domains annotated. Overlapping boxes denote overlapping ORFs.

Searches for deer mouse ERVs in grasshopper mouse, prairie vole, and house mouse suggest that most (9 of 14) deer mouse ERVs arose before the divergence between the deer mouse and its close relative, the grasshopper mouse (∼5-11 MYA) [35,36], but after the divergence between their shared lineage and the prairie vole (∼18 MYA) [37,38], and are thus lineage-specific relative to house mouse (Figure 4B). Additionally, one ERV (Beta_Pman-ERV_cluster-5) evolved even more recently, after the divergence between the deer mouse and grasshopper mouse. Moreover, LTR identity for ERVs in each respective cluster generally concur with these results (Supplementary Table 6). Given the recent origins of deer mouse ERVs and since ERVs frequently arise in new species through the invasion and subsequent endogenization of exogenous retroviruses, we searched for lineage-specific ERVs that display strong homology to a known exogenous retrovirus. We identified one potential case of a recent endogenization of an exogenous Feline Leukemia Virus (FeLV), or closely related virus, in the ancestor of the deer mouse and grasshopper mouse within the last ∼11-18 million years (Figure 4C) [37,38].

### Some deer mouse ERVs may still be infectious

Although ERVs only require *gag* and *pol* genes to mobilize in the germline, ERVs with intact *env* genes can also infect other cells. Given the recent evolution of ERVs in the deer mouse, we inspected all intact ERVs as well as ERV subfamily consensus sequences for intact *env* genes. We find no evidence for *env* genes in mysTR related subfamilies, consistent with previous studies on mysTR [18] (Figure 4D; Supplementary Table 4). However, we find putatively intact *env* genes in multiple other ERV clusters, suggesting that some deer mouse ERVs are still capable of horizontal transmission (Supplementary Table 4, 6). One of these is Gamma_Pman-ERV_cluster-10, consistent with previous observations that other Leukemia Viruses remain infectious [39,40] (Figure 4D). We also observe evidence of an intact *env* gene for ERVs within the Beta_Pman-ERV_cluster-5. Beta_Pman-ERV_cluster-5 ERVs are absent in grasshopper mice and thus represent some of the most recent ERVs to arise in the deer mouse (Figure 4B, D). Beta_Pman-ERV_cluster-5 *env* genes show sequence homology to the *env* IAP elements in house mice, which are also capable of intracellular transmission [41], suggesting possible origin from a similar retrovirus (Supplementary Table 4, 6).

### Negative selection shapes TE distributions

Most TE insertions are deleterious or neutral, and the distribution of TEs in a genome is primarily shaped by negative selection. In the deer mouse genome, TEs are generally well represented in coding genes, accounting for nearly 25% of genic nucleotides, but are relatively absent from coding exons, suggesting strong purifying selection on new insertions in coding exons (Figure 5A). We also observe considerable representation of TEs in long noncoding RNAs (lncRNAs), consistent with observations in other species [42]. Comparison of TE occupancies across chromosomes reveals that ERVs and LINEs exhibit higher occupancies on the X chromosome (occupying ∼15% and ∼17 of the X chromosome respectively, compared to an average of ∼11 and ∼10% for other chromosomes; Figure 5B). This pattern is not observed for SINEs and likely reflects the more frequent removal of longer TEs such as LINEs and ERVs on autosomes by recombination [43,44]. ERV insertions around protein-coding genes are also usually deleterious since ERVs contain complex internal regulatory elements that can disrupt gene expression. Consistent with this, ERVs are generally distant from genes and significantly more distant from genes in the same orientation (Mann Whitney U Test, P<0.0001; Figure 5C). It is worth noting that this bias is most pronounced for mysTR related ERV subfamilies, suggesting that these ERVs are highly deleterious.

**Figure 5:**
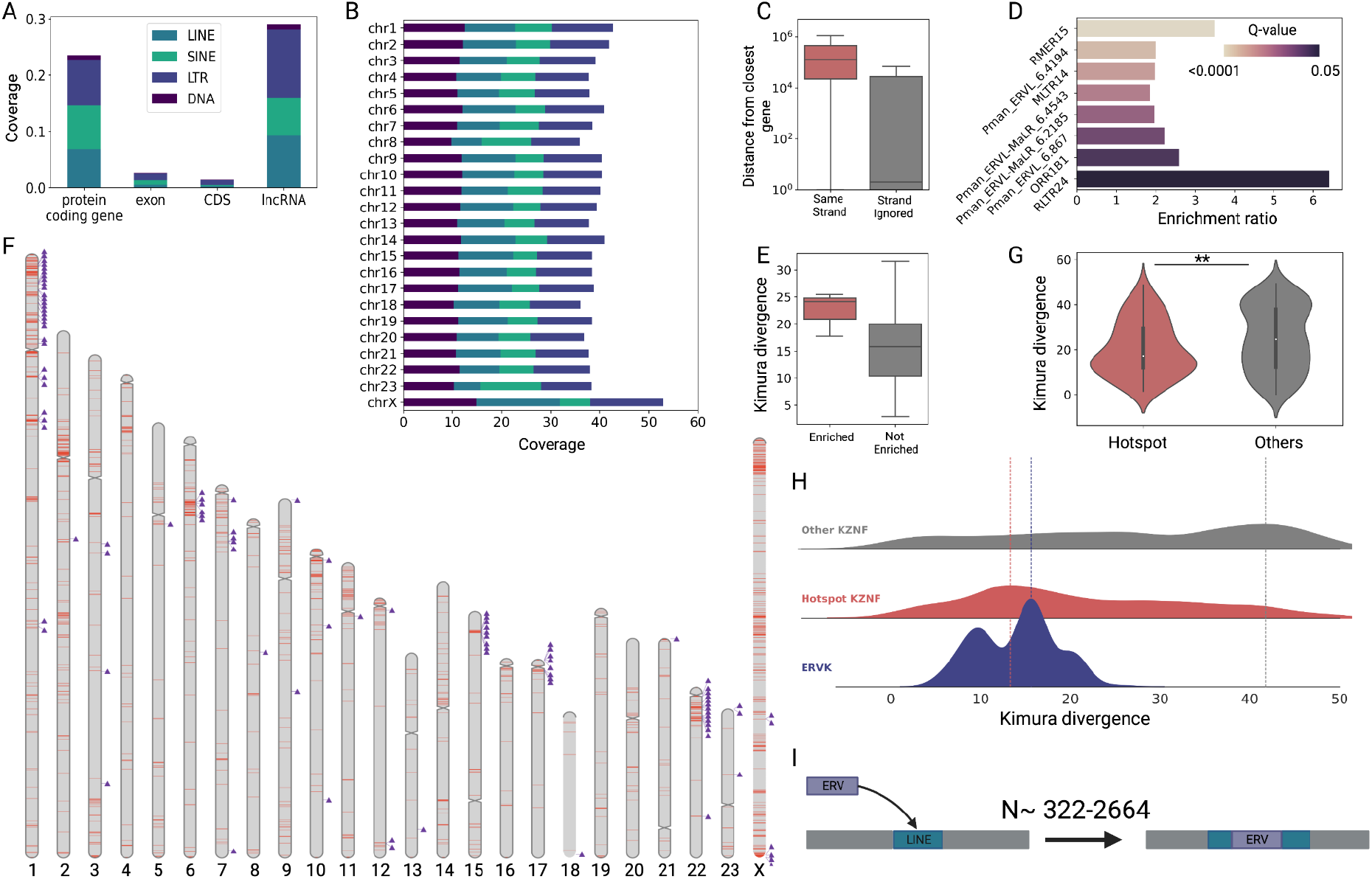
Genomic distribution of TEs in the deer mouse genome. (**A**) Respective coverage for different TE subclasses across genomic features. Coverage is defined as the proportion of nucleotides attributed to TEs for a given feature. CDS = protein coding sequence; lncRNA = long noncoding RNA. (**B**) Respective coverage for TEs across deer mouse chromosomes. (**C**) Box plots showing the distribution of ERV distances from the closest gene on the same strand (red) versus when strand is ignored (gray). (**D**) Enrichment ratio (number observed/expected) and bonferroni-corrected Fisher’s Exact Test P-values (Q-values) for ERV subfamilies enriched within the 5 kb region upstream of genes in the same orientation. (**E**) Within-subfamily CpG corrected Kimura divergence for ERV subfamilies enriched within the 5 kb region upstream of genes in the same orientation (red) compared to all other ERV subfamilies (gray). (**F**) Genomic distribution of ERV hotspots (red) across chromosomes. Lineage-specific KZNF genes are indicated (purple triangles) and are enriched in ERV hotspots. **(G)** KZNFs in ERV hotspots (red) show lower Kimura divergence than other KZNFs (gray), suggesting that they are younger. **(H)** Kernel density estimates for the distribution of Kimura divergences for KZNFs outside ERV hotspots (gray), KZNFs in ERV hotspots (red), and ERVKs (blue). Color-coded dotted lines show the peak value for each distribution. **(I)** Cartoon displaying an ERV insertion interrupting a formerly intact LINE. *N* represents the range of observed candidate instances of ERV-mediated LINE interruption.

### Some ERV subfamilies have possible regulatory function

While most ERV subfamilies show patterns suggesting deleterious effects on neighboring genes, others display patterns consistent with possible regulatory function. ERV LTRs are often co-opted for important regulatory functions over evolutionary time [45]. We find some ERV subfamilies are enriched within the 5-kb region upstream of genes, suggesting selection on these ERVs to minimally affect neighboring gene expression or the possibility of host co-option (Figure 5D). These ERVs also display significantly higher within-subfamily divergence relative to other lineage-specific deer-mouse ERVs (Mann Whitney U Test, P=0.0048), suggesting that they primarily represent inactive subfamilies (Figure 5E). Additionally, some subfamilies, including MT2B1 and ORR1B1, represent lineages of ancestrally shared elements that have been co-opted for regulatory functions in other mammalian species [46]. Functional enrichment tests on neighboring genes for each of these subfamilies yield few enriched categories. Regardless, given the frequent and recurrent lineage-specific ERV co-option events observed across mammals [47–49], these ERVs represent promising candidates for independent co-option events in the deer mouse.

### ERVs accumulate in “hotspots” enriched for Kruppel-associated box domain-containing zinc finger genes

The distribution of ERVs in the genome is largely biased towards specific regions, or “hotspots”, which are enriched in Kruppel-associated box domain-containing zinc finger genes (KZNFs). We define “hotspots” as regions of the genome in the top 95th percentile of ERV density, where ERV density is the proportion of nucleotides attributed to ERVs in a given 100-kb genomic window (Figure 5F). Lineage-specific ERVKs constitute over 70% of ERVs in hotspots, suggesting that these genomic structures are likely also lineage specific. ERV hotspots are largely devoid of genes, and we observe a strong negative correlation between gene density and ERV density overall (GLM, P<0.0001). However, we do observe some genes in ERV hotspots. We performed a gene ontology (GO) enrichment analysis for genes in ERV hotspots and found significant enrichment for one biological process term: “regulation of transcription, DNA-templated” (Fisher’s Exact Test, Q<0.00001). Scrutiny of genes overlapping ERV hotspots that match this GO term reveals that ∼85% (100/118) are deer mouse-specific Kruppel-associated box domain containing zinc finger genes (KZNFs) (Figure 5C). We define deer-mouse specific KZNFs based on refseq’s annotation of genes that do not have orthologs in other species. We find that deer mouse-specific KZNFs specifically are enriched in ERV hotspots, with ∼32% (100/312) of KZNFs occurring in ERV hotspots, despite the fact that ERV hotspots only represent <5% of the genome (Fisher’s Exact Test, P<0.00001).

### Coevolution of ERVs and KZNFs

The primary function of KZNFs is to suppress ERV activity [11,50,51]. KZNF gene clusters evolve rapidly through a birth-death model under positive selection and often expand in response to lineage-specific ERV activity [52–54]. The co-localization of KZNF genes and ERVs in genomic space is curious and has been observed previously in the house mouse [55]. Although this observation could simply be explained by relaxed selection on nonessential KZNF genes, two alternative, non-mutually exclusive hypotheses could explain the observed colocalization between KZNFs and ERVs: (1) KZNFs use neighboring ERVs as regulatory sequences to respond to the global derepression of ERVs [56,57] or (2) an ERVs contribute to KZNF gene family evolution by facilitating rapid gene turnover in these regions. Indeed, in support of the latter, ERVs facilitate structural rearrangements via ectopic recombination, and ERV-rich regions of the genome are often highly plastic. Interestingly, lineage-specific KZNF duplicates in ERV hotspots exhibit significantly lower divergence compared to other KZNFs, suggesting that genes in ERV hotspots duplicated more recently (Mann Whitney U, P=0.0043; Figure 5G). This observation supports a hypothesis in which KZNFs in ERV hotspots duplicate more often, although the evolutionary processes driving this pattern remain unclear.

The relative timing and magnitude of lineage-specific KZNF gene family expansion in ERV hotspots reflects that of lineage-specific ERVK activity. We compared the distribution of candidate gene duplicate divergence for KZNFs in ERV hotspots and KZNFs outside ERV hotspots with the distribution of within-subfamily ERVK divergence (Figure 5H). The distribution of duplicate divergence for KZNFs in ERV hotspots suggests that the largest KZNF expansion occurred just before or around the same time of peak ERVK activity (Figure 5H). Indeed, the median percent divergence for lineage-specific KZNF gene duplicates in ERV hotspots is ∼17.2%, while the within-subfamily divergence for the top three most abundant ERVs in the deer mouse genome is ∼17.4%. This pattern is consistent with a KZNF expansion in ERV hotspots in response to highly active lineage-specific ERVKs. In contrast, the distribution of duplicate divergence for KZNFs not overlapping ERV hotspots shows little evidence for a relationship to lineage-specific ERVK activity (Figure 5H), further suggesting that the observed colocalization between ERVKs and KZNFs may be evolutionarily significant. Furthermore, some KZNF gene clusters display much larger expansions than others: for example, a cluster on chromosome 1 contains >90 genes, representing ∼1/3 of annotated lineage-specific KZNFs in the genome (Figure 5F). This observation suggests that KZNFs in this chromosome 1 cluster, in particular, may play an important role in suppressing ERVs in the deer mouse. We observe another case on chromosome 22, which displays a custer of 48 lineage-specific genes. Since specific KZNF clusters often bind to specific ERV families, the massive invasion of closely related ERVs in the deer mouse predicts expansions of closely related KZNFs [52]. Together, these results suggest that KZNFs in the deer mouse underwent a massive expansion in response to lineage-specific ERV activity.

### Lineage-specific ERV insertions interrupt LINE sequences

In addition to evaluating ERV distributions with respect to genes, we also assessed ERV distributions with respect to other TEs. We were specifically interested in how the observed ERV invasion in the deer mouse might directly impact pre-existing LINEs. ERVs could have directly impacted LINE activity by inserting into and interrupting transposition-competent LINEs. LINE families typically only have from a hundred to a few thousand “master genes” that are transposition-competent in mammalian genomes [5,58–60]. Furthermore, the LINE retrotransposition mechanism is fairly inefficient, and the vast majority of new LINE insertions are defective and incapable of mobilizing thereafter [61,62]. Thus, disruption of many master genes could have a considerable impact on the evolutionary trajectory of LINEs in a species.

To explore the direct impacts of ERV insertions on LINEs, we searched for ERV insertions directly flanked by LINE sequences from the same LINE subfamily. We then filtered for cases in which flanking LINE sequences conjoined at the correct coordinates with respect to the subfamily consensus, forming a full-length LINE. We also initially filtered for LINEs that did not contain any additional TE insertions. These results revealed 322 prospective lineage-specific ERV insertions that interrupt full-length LINEs (Supplementary Table 7). However, this number is likely a considerable underestimate, since it does not include fragmented LINEs that have accumulated multiple indels. If we include fragmented LINEs as well, we find 2664 prospective ERV insertions interrupting LINEs, 900 of which are attributed to the two most abundant ERVK related subfamilies (Pman_ERV2_4.503 and Pman_ERV2_4.24) (Supplementary Table 8). Interestingly, within-subfamily percent divergences for these subfamilies (16.05% and 15.86%) suggests that they invaded just before the decline of LINE gain (see Figure 1B, ∼15%). We postulate that this association is no coincidence. Punctuated LINE interruptions on these scales (322 - >2500 LINE interruptions) would eliminate most functional LINEs in many mammalian species, and even on much smaller scales, could have a catastrophic effect on the evolutionary trajectory of LINEs in a genome, especially given the poor success rates of the LINE retrotransposition mechanism in producing new fully functional LINE copies.

We also explored quantitative patterns of ERV content in LINEs more broadly. To do so, we tested for an enrichment of ERVs in LINEs across ERV subfamilies. Interestingly, we found that several lineage-specific ERVK subfamilies show significant enrichment in LINEs (bonferroni corrected permutation test, α = 0.01, N = 1000; Supplementary Table 9). Pman_ERVK_4.63, for example, interrupts LINEs >46 times more than expected by chance (bonferroni corrected permutation test, Q<0.001). We acknowledge, however, the challenge of identifying the most appropriate null hypothesis for this analysis. Nonetheless, the observed enrichment for lineage-specific ERVK subfamilies, representing 29 of the 36 enriched subfamilies, and absence of enrichment for other ERV subfamilies, is consistent with our hypothesis that ERVK expansion directly impacted autonomous LINE viability (Supplementary Table 9). Together, these results suggest competition between ERVs and LINEs for genomic sites and that ERVKs may have directly impacted the evolutionary trajectory of LINEs in the deer mouse.

### An “ecology of the genome” model for the evolution of the deer mouse genome

In the same way that species compete for space and resources, TEs compete with each other for sites in the genome as well as metabolic resources [9]. TEs can occupy specific niches, which can allow them to coexist with limited competition, but TEs that occupy similar niches are more likely to compete and thereby drive one or another to extinction [9,10]. Furthermore, the relative success of a given TE also depends on host suppression mechanisms and their targets. For example, differential host targeting between two TE families in direct competition could limit the success of one family that would, in the absence of host defense mechanisms, be more fit than the other [10]. Also, because TEs could threaten to kill their host in the absence of host-mediated suppression, it can be advantageous (for both the host and TEs) that host defenses evolve to suppress TE activity [10].

Our model for the evolution of the unique TE landscape observed in the deer mouse genome lends itself to the ideas explained above: the “ecology of the genome” [9]. We postulate that the introduction of mysTR-related ERVs sparked a shift in the deer mouse TE landscape through the following processes (Figure 6): first, mysTR ERVs evaded host defenses upon immigration, which allowed them to expand to large numbers. This hypothesis is supported by the observation that mysTR ERVs are highly divergent from other known retroviruses as well as the remarkable expansion of deer mouse-specific KZNFs following peak ERV activity [18]. In mammals, ERVs are the primary targets of KZNF suppression, whereas LINEs and SINEs are less frequently targeted, probably because ERV insertions are more deleterious [63,64]. These host defenses keep ERVs in check, despite evidence that LINEs and ERVs occupy overlapping niches. In fact, several lines of evidence suggest that LINEs and ERVs are in direct competition. First, many ERVs and LINEs both preferentially integrate into AT rich regions [65–68]. Thus, ERVs and LINEs often inhabit similar regions of the genome and frequently insert within each other [65]. Second, ERV insertions in LINEs (or vice versa) are likely invisible to selection and exhibit a higher rate of fixation relative to deleterious insertions [65]. Under these circumstances, in the absence of host defense mechanisms, we expect the primary driver of ERV or LINE success in the genome to be relative rates of gain (of transposition-competent copies). Thus, we postulate that the massive expansion of mysTR ERVs nearly drove LINEs to extinction in the deer mouse genome.

**Figure 6:**
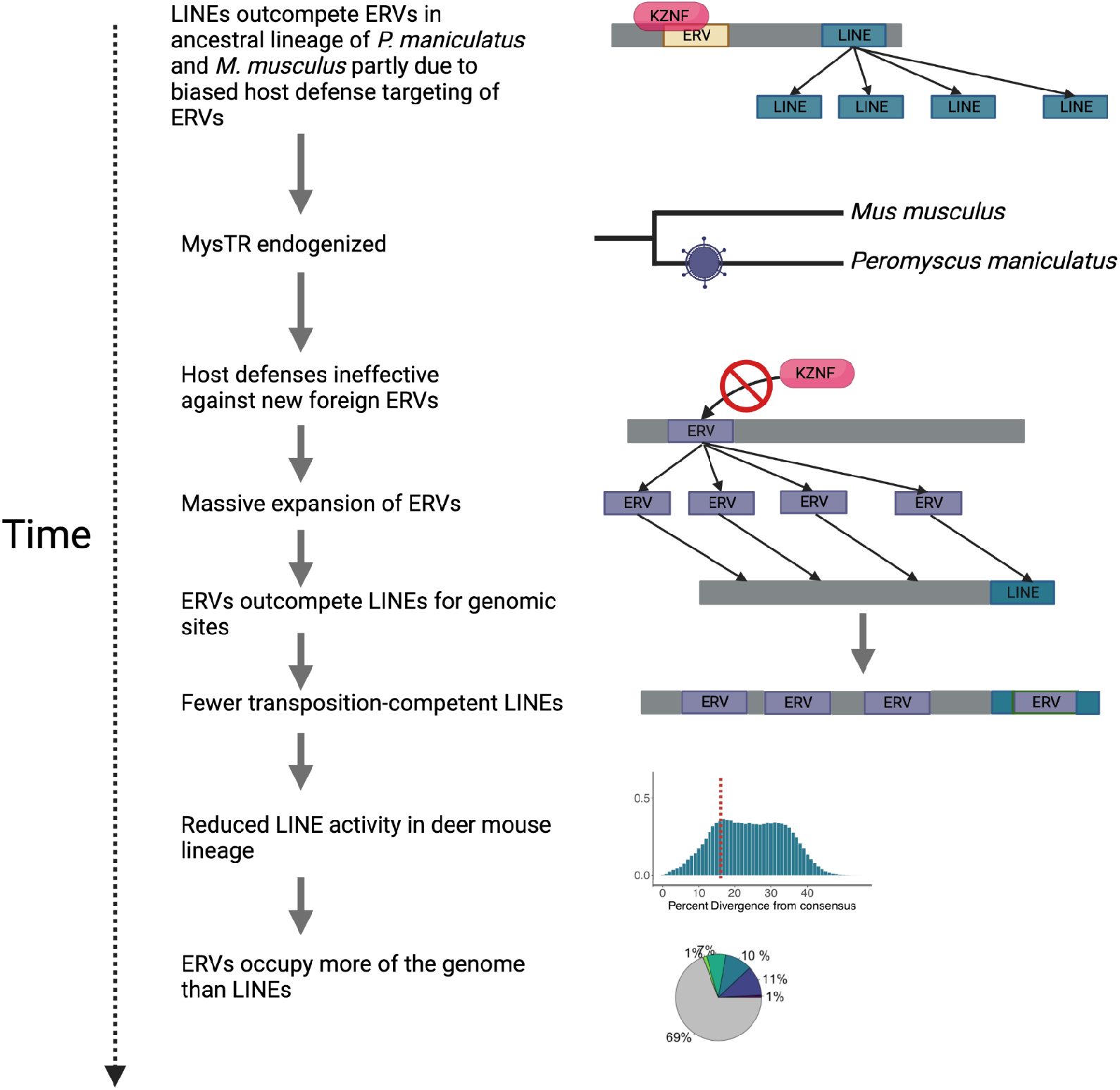
Model for deer mouse genome evolution.

More generally, we propose that this model may explain the loss of LINE activity in other mammals. A subclade of sigmodontine rodents for example (∼13-18 MYA diverged from the deer mouse [36–38,69]), represents one of the few mammalian lineages to have experienced LINE extinction [70]. Consistent with our model, previous studies suggest that LINE extinction in this group followed an invasion of mysTR related ERVs on a similar or possibly larger scale to that observed for the deer mouse [18]. At present, the lack of genome assemblies for sigmodontine rodents makes it challenging to study TEs in these species. Future studies in these and other species that show unique patterns of mammalian genome composition will shed further light on evolutionary conflicts that drive mammalian genome evolution.

### Concluding remarks

Although TE landscapes differ drastically across species, most mammalian genomes are similar and dominated by LINE retrotransposons. The deer mouse, *Peromyscus maniculatus*, represents one of the few exceptions to this pattern: LTR elements occupy more of the genome than LINEs. We find that the unique genomic landscape of TEs in the deer mouse reflects a massive expansion of ERVs as well as a dearth of LINE activity on recent timescales, and that the two are likely associated. Our results demonstrate a broad diversity of ERVs in the deer mouse genome and that the immigration of one particular divergent ERV, mysTR, played an integral role in establishing its unique TE landscape.

Based on these findings, we postulate that the propensity for a mammalian genome to undergo a shift in TE content and/or experience LINE extinction is directly related to its susceptibility to invasion by divergent TEs (in this case, ERVs). Furthermore, the accumulation of ERVs in specific hotspots raises additional questions about how TE-dense regions can affect mammalian genome evolution. Indeed, we would expect such regions to experience structural rearrangements more often than other regions of the genome. Previous studies in the deer mouse have identified many massive inversions (>1 Mb in length), which are polymorphic, even within populations [71,72]. Could these ERV hotspots have played a role in facilitating deer mouse inversions? We also observe enrichment of KZNF gene families which evolve rapidly via duplication under positive selection in ERV hotspots. Is the colocalization of KZNFs and ERVs advantageous for the host due to the increased propensity for KZNF gene family expansion? We show that KZNFs in ERV hotspots are indeed younger than other KZNFs, providing some support for the coevolution of these elements. However, within-population studies are critical to further elucidate this coevolutionary relationship. Together, our results have broad implications and open up a range of opportunities to investigate the evolutionary processes that give rise to the evolution of mammalian genome structure.

## Methods

### Obtaining relevant genomic data

We downloaded publicly available TE annotations for human, *Homo sapiens* (GCF_000001405.40; genome contig N50 = 57,879,411; contig L50 = 18); house mouse, *Mus musculus* (GCF_000001635.27; genome contig N50 = 59,462,871; contig L50 = 15); Norway rat, *Rattus norvegicus* (GCF_015227675.2; genome contig N50 = 29,198,295; contig L50 = 27); and prairie vole, *Microtus ochrogaster* (GCF_000317375.1; genome contig N50 = 21,250; contig L50 = 29,205) from RepeatMaster (http://www.repeatmasker.org/genomicDatasets/RMGenomicDatasets.html) and for grasshopper mouse, *Onychomys torridus* from NCBI (GCF_903995425.1; genome contig N50 = 2,276,141; contig L50 = 308) respectively. We used the deer mouse, *Peromyscus maniculatus*, genome assembly available through NCBI (refseq GCF_003704035.1; contig N50 = 30,111; contig L50 = 23,323) for all genomic analyses. Retroviral sequences for ERV phylogenetic analysis were downloaded from NCBI. Genbank accession numbers and for these sequences are shown in Figure 4A.

### TE discovery and annotation

We used a combination of systematic and manual techniques to identify and annotate TEs in the deer mouse genome. We started by using an approach similar to the EarlGrey pipeline (github.com/TobyBaril/EarlGrey/) [73]. We first identified known rodent TEs in the deer mouse genome using RepeatMasker (version 4.1.2) (https://www.repeatmasker.org/) with a curated set of rodent TEs from the DFAM database [20] and the flags *−nolow, -norna* and *−s*. Next, we constructed a *de novo* repeat library using RepeatModeler2 (version 2.0.1), with RECON (version 1.08) and RepeatScout (version 1.0.5) [74–76]. Maximum-length consensus sequences were generated for putative *de novo* TEs identified by RepeatModeler using an automated version of the “BLAST, Extract, Extend” process through EarlGray [21]. Briefly, EarlGray first performs a BLASTn search to obtain up to the top hits for each TE subfamily [77]. Then, it aligns the 1000 base pairs of flanking retrieved sequences using MAFFT (version 7.453) [78]. Following this, alignments are trimmed using trimAl (version 1.4) with the options (*−gt* 0.6 *-cons* 60) [79]. Finally, consensus sequences are updated using EMBOSS cons (*−plurality* 3) [80]. This process is then repeated five times. Following this, we performed blastx [77] searches against all known deer mouse proteins with parameters (*-max_target_seqs* 25 *-culling_limit* 2 *-evalue* 10e-10) and filtered all TEs with unknown classifications that shared homology with proteins.

Following the automated processes described above, alignments for TE families were individually inspected using AliView [81] and poorly represented positions were manually trimmed as recommended by [82]. Families were also manually realigned using extract_align.py [21] and MAFFT (version 7.453) [78] and then reexamined. Manually curated TE families were then re-clustered using cd-hit-est [83] and families were merged based on the 80-80-80 rule criterion [84]. We also used TE-Aid (https://github.com/clemgoub/TE-Aid) to identify TE-associated ORFs and sequence features such as long terminal repeats when classifying TEs. We combined our final *de novo* TE library with the Rodent DFAM TE library [20] and annotated TEs in the deer mouse genome using RepeatMasker (version 4.1.2) (https://www.repeatmasker.org/). To identify full-length LTR elements, we used LTR_FINDER [85] and LTRharvest [86] through EDTA_raw with the flag *-type ltr* (version 2.0.0) [87], which also report LTR divergence for each element. TE annotations were defragmented and refined using RepeatCraft with the flag *−loose* [88], and overlapping annotations were resolved in favor of the longer element using MGKit (version 0.4.1) filter-gff [89].

### Identifying functional machinery for putatively autonomous TEs

To identify the protein machinery of potentially autonomous LINE and LTR elements, we extracted all LINE elements longer than 2700bp and LTR elements longer than 5000bp from the deer mouse genome. Then, we also used TE-Aid (https://github.com/clemgoub/TE-Aid) to identify ORFs in each retrieved LTR and LINE element with homology to known TE genes. We used hmmer [32] and relevant hmms available from GyDB [90] and PFAM [91] to identify retroviral protein domains as well as NCBI’s conserved domain search tool [92,93].

### Calculating *k* coefficients

We calculated the DNA loss coefficient *k* [24], using the formula *E = A e-kt*, where *E* is the amount of extant ancestral DNA in the species considered, *A* is the ancestral assembly size, and *t* is time. We calculated *E* for each species by subtracting the amount of genomic DNA attributed lineage-specific TEs from the amount of DNA attributed to ancient shared mammalian TEs (retrieved from [94]). We used 2.8Gb for *A* and 100 million years for *t* as in [94].

### Identifying nonautonomous ERV-like elements

Since the deer mouse genome was produced primarily with short reads, most ERVs have internal gaps or strings of low quality or ambiguous nucleotides. Thus, to decipher nonautonomous ERV-like elements from autonomous ERVs, we used a strict criterion. For a given ERV subfamily to be considered nonautonomous, we required at least five full-length copies which lack identifiable ORFs as well as ambiguous nucleotides. We performed global pairwise alignments between nonautonomous and autonomous ERVK consensus sequences using AVID with default parameters [95]. We visualized alignments using VISTA [96].

### ERV classification and phylogenetic analysis

We used two complementary approaches to classify deer mouse ERVs. First, we examined e-value statistics in the output from our GyDB hmm scans to discern which viral reverse transcriptase domain hmm best fit each ERV. In addition, we also used a phylogenetic approach. We annotated ERVs with their viral origin as predicted by our hmm scans. Next, we downloaded several endogenous and exogenous retroviruses from NCBI (accessions shown in Figure 4A), extracted their RT domains and annotated them with their respective viral clade. Then, we filtered sequences with large strings of ambiguous characters, performed a multiple sequence alignment of RT genes using MAFFT [78], and generated a maximum likelihood based phylogeny using IQ-TREE [97] with a GTR+G model (general time reversible model with unequal rates and unequal base frequencies and discrete gamma rate heterogeneity) [98]. We analyzed and edited the resulting phylogeny using ete3 (version 3.1.2) [99], collapsing clusters of deer mouse ERVs into representative nodes. We visualized the phylogenetic tree using the Interactive Tree Of Life (ITOL) [100] and FigTree (version 1.4.4) (https://github.com/rambaut/figtree).

### Searching for deer mouse ERVs in other species

To search for deer mouse ERVs in the house mouse, prairie vole and grasshopper mouse genomes, we performed local BLASTn [77] queries for each full-length deer mouse ERV to each respective genome. We ran BLASTn [77] with the flag *-outfmt 6* and required a minimum alignment length of 400bp and minimum percent identity of 75 to limit possible erroneous hits. As a proof of concept, we also made sure our results were consistent with expectations based on LTR divergences. For example, we would not expect an ERV with highly divergent LTRs (a signature of a more ancient insertion) to be specific to the deer mouse. We also performed broader BLASTn queries against NCBI’s nucleotide database. Queries for Gamma_Pman-ERV_cluster-10 sequences only yielded high-confidence hits in deer mouse species, the grasshopper mouse, and the Feline Leukemia Virus (FeLV) reference genome. A neighbor-joining phylogeny constructed from deer mouse Gamma_Pman-ERV_cluster-10 sequences, homologous ERVs in the grasshopper mouse genome, and the FeLV reference suggests a scenario in which Gamma_Pman-ERV_cluster-10 originated in the common ancestor the deer mouse and grasshopper mouse from FeLV or another closely related exogenous virus between 11 and 18 million years ago [36].

### TE distribution analysis

We used bedtools intersect [101] to find overlaps between TE annotations and gene feature annotations. We used bedtools closest [101] with the parameter *-s* to identify TE distances from the nearest gene on the same strand and again with default parameters to ignore strand. All functional enrichment tests were performed using goatools [102]. We also tested for enrichment or depletion of TEs 5 kb upstream of genes in the same orientation. Specifically, for each TE subfamily, we randomized all TE locations on each chromosome and compared the number of TEs within 5 kb of genes upstream in the same orientation with the observed value. We repeated this 10,000 times to obtain a P-value. Functional enrichment Fisher’s Exact P-values and permutation test P-values were adjusted using the bonferroni method to obtain Q-values. We used bedtools coverage [101] to calculate ERV and gene density along 100 kb windows in the genome. ERV hotspots were defined as windows which exhibit ERV densities within the top 95th percentile. Figure 5F was produced using RIdeogram [103].

### KZNF gene family analysis

We defined deer mouse-specific zinc finger (ZF) genes as genes which do not have recognizable orthologs as annotated by NCBI. We employed hmmscan [32] using KRAB hmms downloaded from PFAM [91] to identify KRAB domain-containing ZFs (KZNF). Then, we performed a multiple sequence alignment of all KZNFs using Clustal omega [104] with the parameters *--use-kimura* and *--full* in order to simultaneously produce a pairwise Kimura divergence matrix across all genes. We constructed a subsequent phylogeny using IQ-TREE [97] with a general time reversible model. To test for phylogenetic clustering of KZNF that overlapped ERV hotspots, we used phyloclust through RRphylo R package [105] with 100 simulations. Since KZNF genes evolve via a birth-death process, we define duplicate genes as genes that exhibit the lowest divergence among all pairwise comparisons.

### ERV-mediated LINE interruption

To identify candidate LINEs interrupted by ERVs, we searched for LINE fragments which would be full length (>5000bp) if connected but exhibit an ERV sequence which splits them with respect to their subfamily consensus (Supplementary Table 7). This yielded 322 candidate ERV-mediated LINE interruptions, 121 of which represented lineage-specific LINEs. In this first analysis, we excluded LINEs which showed more than two fragments. If we include those as well, we find 2664 candidate ERV-mediated LINE interruptions. We employed a permutation test to quantitatively assess biased representation of ERVs in LINEs. We did this separately for each ERV subfamily. To do this, we compared the observed number of ERV insertions inside LINEs (ERV sequences flanked on both sides by LINE sequences from the same subfamily) to expectations by randomization 1000 times. We calculated the proportion of iterations that ERVs interrupted LINEs more than expected to obtain a P-value for each ERV subfamily. Then we performed a bonferroni correction to obtain Q-values (Supplementary Table 8).

## Acknowledgments

The authors thank Daniel Hartl, Russell Corbett-Detig, Andreas Kautt, Scott W. Roy, and Timothy B. Sackton for helpful discussions and feedback on this manuscript. HEH is an Investigator of the Howard Hughes Medical Institute.

